# Open Dataset of Developmental Timelines for Corresponding Milestones in Mice, Rats, and Humans

**DOI:** 10.64898/2026.10.09.757751

**Authors:** Eugenia Amici, Rafael Grimson, Pablo Vázquez-Borsetti

## Abstract

Translating developmental time across species is essential for the appropriate design and interpretation of preclinical neurodevelopmental research, yet no open, unified resource currently integrates high-quality developmental milestone data for mouse, rat, and human. Here we present a curated, open-access dataset of general and neurodevelopmental milestones spanning these three species, expanding a previously published rat–human database with newly compiled mouse data. Following a systematic literature search, 783 records covering 371 unique developmental milestones were extracted, standardized to days after fertilization, and validated through a structured curation and quality-control process. Exploratory analyses indicated that the developmental relationship between humans and rodents is markedly non-linear, with milestones becoming increasingly dispersed as development progresses. The observed pattern is consistent with the quarter-power model with an additive constant identified in a companion modeling study. Accordingly, applying a quarter-power transformation to human DAF substantially improved the linearity of developmental trajectories across species. Comparison of body and brain milestone clusters showed no detectable differences in scaling relative to humans, in line with similar underlying rules for somatic and neural development between rodents and humans. The dataset, together with its documented curation methodology, provides a transparent and extensible resource for cross-species developmental translation and a useful tool for informing the generation of animal models.

## Introduction

Animal models are indispensable in neuroscience and biomedical research, enabling the controlled and ethical exploration of biological mechanisms underlying human health and disease. Among these, rodents such as mice and rats are the most widely used species due to their genetic, anatomical, and physiological similarities to humans, combined with practical advantages such as their small size, short generation times, and extensive genetic tractability, see (Bryda 2013). However, translating findings from developmental studies in these species to humans remains challenging because developmental processes unfold over markedly different timescales and follow non-linear trajectories across species (Campos Eusebi et al. 2024; Otis and Brent 1954).

Across early comparative developmental studies, rats were found to be less developmentally mature at birth than humans. This difference was evident from the observation that many developmental milestones occurring prenatally in humans take place postnatally in rats, with many of these observations being well documented in the developing brain (Dobbing and Sands 1979; Donaldson 1918; Romijn et al. 1991).

This developmental disparity posed a significant challenge for modeling perinatal insults and led to the proposition that the rat brain reaches a developmental stage comparable to that of a human newborn only around postnatal days 7–12, an approach that was questioned by previous research (Campos Eusebi et al. 2024). Subsequent evolutionary and comparative studies showed that developmental trajectories across species tend to be non-linear (Clancy et al. 2001). In particular, logarithmic transformations of developmental age were found to reduce the convexity observed when developmental stages were compared across species. This approach was subsequently applied to the rat–human comparison by our group (Campos Eusebi et al. 2024), in which we also compiled an open-access dataset of rat neurodevelopmental milestones by systematically curating empirical data from the literature. This work demonstrated that logarithmic models significantly outperformed linear models in describing cross-species developmental relationships. However, logarithmic transformations failed to fully linearize the data, indicating that the relationship between rat and human developmental ages requires an alternative mathematical formulation. Importantly, this work also addresses a prevalent misconception in comparative biology. Translating ages between species cannot be reduced to a direct equivalence. Instead, this relationship is more accurately modeled as a range of corresponding developmental periods, as individual milestones are not perfectly aligned across species and progressively diverge following fertilization.

For mice and humans, the foundational embryological work of Otis and Brent deserves particular mention (Otis and Brent 1954). They compared morphological landmarks across the two species, yet few studies have subsequently addressed this species pair comprehensively. Notably, while Otis and Brent relied on human bibliographic data, the mouse data were generated ad hoc under controlled conditions. Although pioneering, this methodological asymmetry introduces an important limitation: the low variability inherent in a single-source dataset may limit the generalizability of the inferred temporal relationships. Consequently, datasets aggregated from multiple independent sources are needed to provide more robust and representative estimates of comparative developmental timing.

Despite these historical advances, the field still lacks a unified, open-access resource integrating high-quality, curated general and neurodevelopmental milestones across the three species most commonly used in translational neuroscience: mouse, rat, and human. Existing comparisons remain scattered across publications, rely on heterogeneous criteria, or are restricted to single species pairs. The absence of a comprehensive, transparent database hinders rigorous assessments of temporal equivalence, limits reproducibility, and constrains the development of computational approaches in developmental biology.

To address this gap, we present a novel, curated comparative dataset of neurodevelopmental milestones encompassing mouse, rat, and human. Expanding upon our previous rat-human dataset (Campos Eusebi et al. 2024), this resource incorporates newly curated mouse developmental data. As before, the data were extracted via a systematic review of the empirical literature, encompassing structural, functional, and molecular events across multiple brain and body regions and developmental processes. Each entry is annotated with the event description, anatomical location, species, age, and the original bibliographic source, thereby enabling flexible filtering and cross-species alignment. The dataset is publicly available and version-controlled to encourage community contributions and iterative refinement. In a companion paper, we evaluated alternative mathematical models for translating developmental time across mouse, rat, and human. Among the models tested, a quarter-power model with an additive constant provided the best overall performance, capturing the nonlinear relationship between developmental age across species and providing a quantitative framework for estimating equivalent developmental ages. The present dataset provides the empirical foundation for these modeling efforts, while the companion study focuses on the development and evaluation of quantitative approaches for cross-species developmental translation. Here, we detail the curation methodology, the dataset structure, and initial analyses demonstrating its utility for translating developmental time. Ultimately, this work aims to strengthen the translational bridge between preclinical neurodevelopmental research and human biology, fostering more accurate experimental design and data interpretation.

## Methods

### Study design

This study was based on the construction and curation of an open dataset of neurodevelopmental and general developmental milestones in humans, mice, and rats, obtained through the systematic integration of previously published quantitative information from the scientific literature. The data were complemented with previously published comparative developmental datasets of rat and human from both our group and external sources. From this combined dataset, comparative analyses and mathematical modeling were performed to describe the temporal relationships in neurobiological development across species. Given its complexity, the mathematical modeling is presented separately in the companion paper. The study adopted a retrospective and quantitative approach, combining the critical collection of empirical data with statistical analysis and nonlinear regression techniques to build a model of temporal equivalence among mouse, rat, and human development.

### Search strategy and study selection

The literature search was conducted systematically across international scientific databases, including PubMed and Google Scholar. Initially, combinations of keywords related to neurodevelopment were considered. However, because the search patterns were highly complex, a comprehensive list of keywords would have been substantially larger than the set of developmental milestones analyzed. Therefore, a list of developmental milestones was generated, and each milestone was subsequently searched across the three species: humans, rats, and mice. The search was limited to articles published in English.

Studies were included if they met the following criteria:

1. Reported original empirical data on neurodevelopmental milestones in humans, rats, and/or mice.
2. Provided explicit age information for the reported developmental milestone. 3. Were published in peer-reviewed journals. 4. For datasets compiled by other authors, individual records were curated according to the same criteria, and records that did not meet these criteria were excluded.

Studies or records were excluded if they met any of the following criteria:

1. Did not provide quantifiable information on developmental milestones. 2. Were opinion articles, reviews, or secondary studies that did not correspond to original data sources or, in the case of compiled datasets, did not provide direct citations to the original sources. 3. Contained information inconsistent with more robust or better-supported sources.

References were managed using Zotero. Each selected article underwent an initial critical appraisal to assess its relevance and methodological quality prior to data extraction.

A “developmental milestone” was defined as any temporally discretizable developmental event related to cerebral or bodily processes that could be associated with an explicitly reported age.

### Data extraction, standardization and organization

Relevant data were manually extracted and organized into a structured dataset. For each record, the following variables were collected (column names in parentheses): species (’Species’), developmental milestone (’Parameter’), age (’DAF’), methodological notes (’Obs’), bibliographic reference (’Reference’), DOI, cluster membership (’Cluster’, categorized as brain or body), gestational age (’GA(H+14)’), and a traceability field (’Quote’) containing relevant textual excerpts and/or associated figures or tables supporting the data. An overview of the dataset variables is provided in the data dictionary (Table 1), serving as a quick reference to their types, and definitions. When multiple studies reported different ages for the same milestone and species, independent records were retained and, subsequently, the mean was used as a representative estimate. The mean was preferred over the median whenever both were reported. Ages were standardized as days after fertilization (DAF), with fertilization defined as day 0. Unless otherwise specified in the original studies, embryonic day 1 (E1), gestational day 1 (GD1), and post conception day 1 (PCD1) were considered equivalent for the purposes of age standardization. For humans, a mean gestational duration of 282.2 days was obtained from the literature (Bergsjø et al. 1990; Gjerdevik et al. 2026), corresponding to 268.2 days from fertilization to birth. For rats, a gestational duration of 22.5 days was used (Hain 1936), whereas a gestational duration of 20 days was used for mice (Dewar 1968). For studies reporting postnatal ages, birth was defined as postnatal day 0 (PND0), and postnatal ages were converted to DAF by adding the corresponding prenatal duration from fertilization to birth to the reported PND. When ages were reported in years, a conversion of 365.25 days per year was used. Ages reported in months were converted using 30.44 days per month.

**Table 1:** Data dictionary for the developmental dataset.

| Column | Type | Unique values | Description |
| --- | --- | --- | --- |
| **Species** | string (categorical) | 3 | Species the observation belongs to. Values: **Human** (377), **Mouse** (206), **Rat** (201). |
| **Parameter** | string (categorical) | 371 | Name/description of the developmental milestone (e.g., “GE volume peak”, “Birth”, “Eye opening”). Curated/standardized across sources to avoid duplicate naming of the same event. |
| **DAF** | float | 216 | **Days After Fertilization** — standardized age at which the milestone occurs, with fertilization defined as day 0. This is the primary age variable used throughout the analysis. When multiple sources report different ages for the same Species+Parameter, records are kept separately (not averaged in the raw file); the mean is computed downstream as the representative value. |
| **Reference** | string | 391 | Bibliographic citation (author, year) for the source of the record. |
| **Obs** | string | 108 | Free-text curator notes, e.g., justification for excluding or adjusting a value, discrepancies with the cited source, or data-quality caveats. Populated only when a note was needed (missing in 621 rows). |
| **Quote** | string | 412 | Traceability field: verbatim excerpt from the source text and/or a reference to a supporting figure or table supporting the reported value. |
| **DOI** | string | 403 | Digital Object Identifier of the reference, used as a unique key for reference-level aggregation (e.g., counting records per source). Missing for a few older/non-indexed sources (3 rows; see **Obs/link** for alternate access). |
| **link** | string | 22 | Alternative URL to the source when no DOI is available (e.g., HathiTrust book scans). Only populated for non-DOI sources (missing in 694 rows). |
| **Cluster** | string (categorical) | 2 | Tissue/domain classification of the milestone: **Brain** (308) or **Body** (472). Missing only for the 4 **Birth** reference rows, which are used to anchor the per-species birth date. |
| **Data_curator** | string (categorical) | 8 | Name of the person/team who curated the record, e.g., **Otis 1954**, **Ohmura 2017**, **Campos-Iorii 2023**, **Amici, Cottam 2024**, **Godlewski 1997**. Used for provenance and to audit curation batches. |
| **GA(H+14)** | float | 47 | Human **Gestational Age** counted from the last menstrual period (LMP), computed as **DAF + 14**. The 14-day offset represents the assumed interval between LMP and fertilization. Only meaningful/populated for **Species == Human**; provided as a convenience for reporting ages in the clinically conventional GA scale alongside DAF (populated in only 58 rows total). |

In cases where studies reported narrow age ranges, the midpoint was used as the representative value. For wider age ranges or events explicitly described as having a start and end, the corresponding values were recorded separately as “start” and “end”.

Data organization was performed using spreadsheets and online collaborative tools, enabling version control and continuous database updates.

### Data Curation

The curation process involved a systematic review and manual validation of each entry included in the dataset by a researcher with expertise in the field. In addition, the terminology used to describe developmental milestones was standardized to ensure comparability across species and to avoid duplication of the same milestone under different names. The inclusion and exclusion criteria, as well as any adjustments made to the original data, were explicitly documented. Entries that could not be traced back to their original sources were excluded from the dataset, even if they had been previously curated by other authors. The curated dataset was then subjected to a quality-control process that included cross-checking entries against the original sources and verifying age standardization and consistency across species. Outliers were identified and flagged for further review. Outliers were removed only when the original study was found to be inconsistent with more robust sources or when there was a documented reason to consider the reported value unreliable. The data were compiled and curated entirely manually, without the use of AI assistance.

### Statistical Analysis and Modeling

The quantitative analysis and the validation of the mathematical models are detailed in the companion paper. The present study focuses on complementary analyses, beginning with an exploratory data analysis (EDA) of the dataset. For the statistical modeling, Ordinary Least Squares (OLS) regression was applied in cases where no evident heteroscedasticity was observed, such as in the developmental relationship between the rat and the mouse. Conversely, weighted regression models were utilized when heteroscedasticity was detected.

When heteroscedasticity was detected, weights for the weighted least squares (WLS) fits were estimated using a two-stage procedure. An initial OLS regression was fitted to obtain preliminary residuals, and the corresponding squared residuals were computed. A variance model was then fitted by regressing the logarithm of the squared residuals on the logarithm of the independent variable, and the fitted values were exponentiated to obtain an estimate of the conditional variance for each observation. Weights were defined as the reciprocal of these estimated variances, so that observations associated with lower estimated variance were assigned proportionally greater weight in the subsequent WLS fit. This data-driven procedure allowed the variance to be modeled as a smooth function of the predictor, correcting for heteroscedasticity without imposing a fixed parametric weighting scheme.

In certain instances, Locally Estimated Scatterplot Smoothing (LOESS) was utilized as a non-parametric smoothing technique. This approach allowed for a visual and quantitative comparison of the data trends against the fit provided by our mathematical model.

All analyses were conducted in Python. NumPy and Pandas were used for data manipulation; SciPy and Statsmodels for statistical analyses and regression modeling; and Scikit-learn for model validation. Data visualization was performed using Matplotlib and Seaborn. The working environment utilized was Visual Studio Code (VS Code) with AI-assisted support (Copilot).

## Results

### Dataset composition and distribution

In this work, we present the assembly of a dataset containing developmental data from laboratory rodents. The curated dataset comprised 783 records covering 371 unique developmental milestones across the three species (Figure 1). Within this dataset, 47 duplicated rows (based on Parameter + Species) were identified, corresponding to 22 unique parameter and species combinations. Observations were distributed across two broad tissue categories, with the majority corresponding to Body-related milestones and the remainder to Brain-related milestones (Figure 1a). Regarding species composition, Human contributed the largest share of records as most milestones were paired between rodents and humans (Figure 1b). Specifically, the detailed overlap profile of the 371 unique milestones revealed that 179 were shared exclusively between Human and Mouse, 167 between Human and Rat, 9 between Rat and Mouse, and 16 were common to all three species. Most observations originated from previously published or externally curated sources (87.7%, n = 687), while the remaining 12.3% (n = 96) corresponded to milestones specifically compiled for the present study (Figure 1c). Across all three species, the majority of records described prenatal milestones (Figure 1d–f).

**Figure 1:**
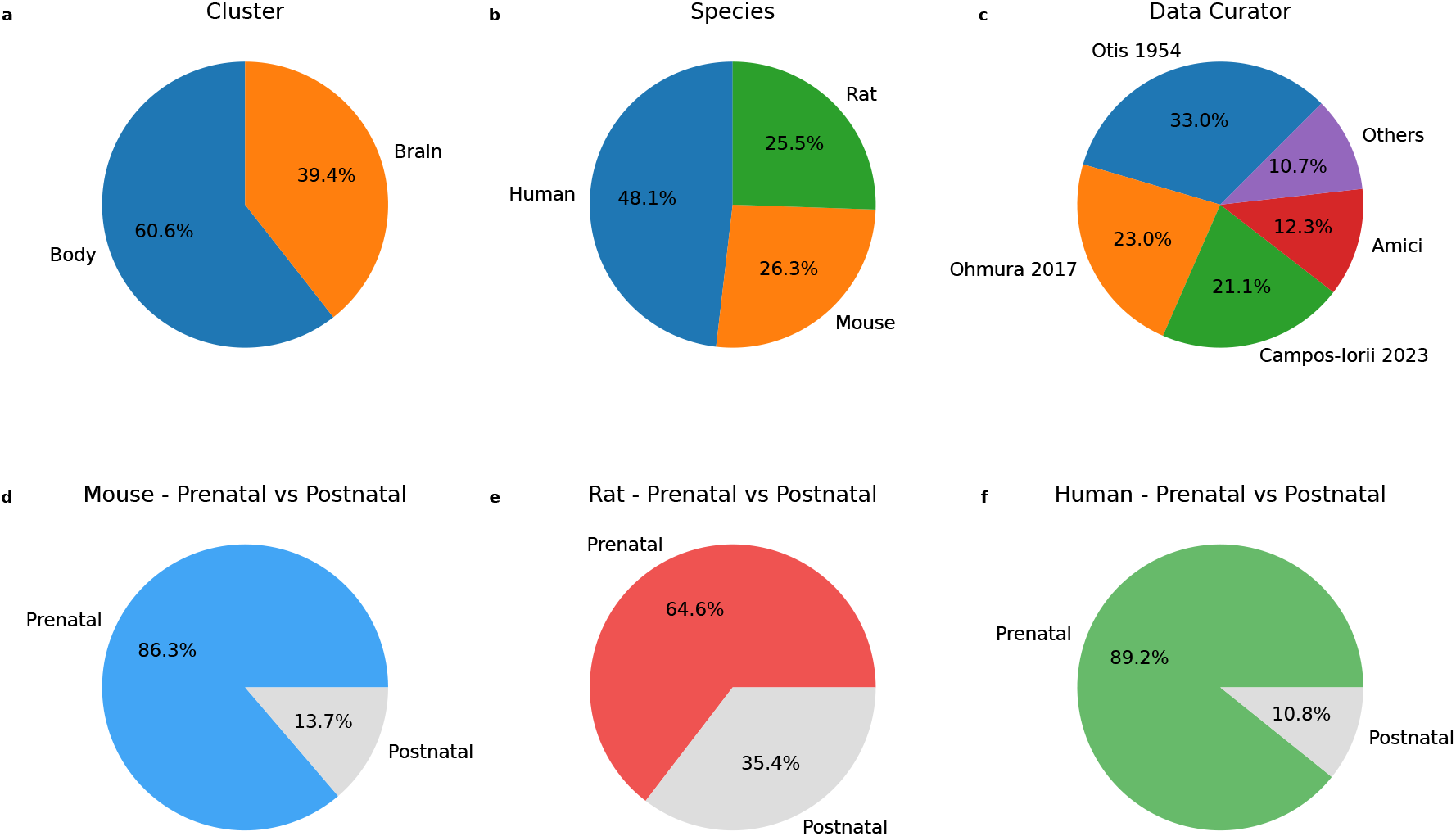
Overview of dataset composition and distribution across tissues, species, data sources, and developmental stages. The dataset comprises measurements from two major tissue categories, Body and Brain (a). Distribution of observations among species, showing a larger contribution from Human data and relatively balanced contributions from Mouse and Rat (b). Distribution of observations according to data source and curator, distinguishing between external datasets and data compiled in-house. Data curated by Amici represent observations specifically compiled for the present study, whereas the remaining observations were obtained from previously published or externally curated datasets (c). Overall distribution of observations across prenatal and postnatal developmental stages. Developmental-stage composition within each species, showing the relative contribution of prenatal and postnatal observations for Mouse (d), Rat (e), and Human (f).

The repetition frequency of digital object identifiers (DOIs) was analyzed to assess the diversity of bibliographic sources used in the assembly of the dataset (Figure 2a). The distribution shows that the vast majority of references (75.7%) are cited only once, 15.6% are repeated twice, 4.2% three times, and only 4.5% appear four or more times. This indicates that the dataset is supported by a broad and diverse literature base, minimizing bias towards isolated studies.

**Figure 2:**
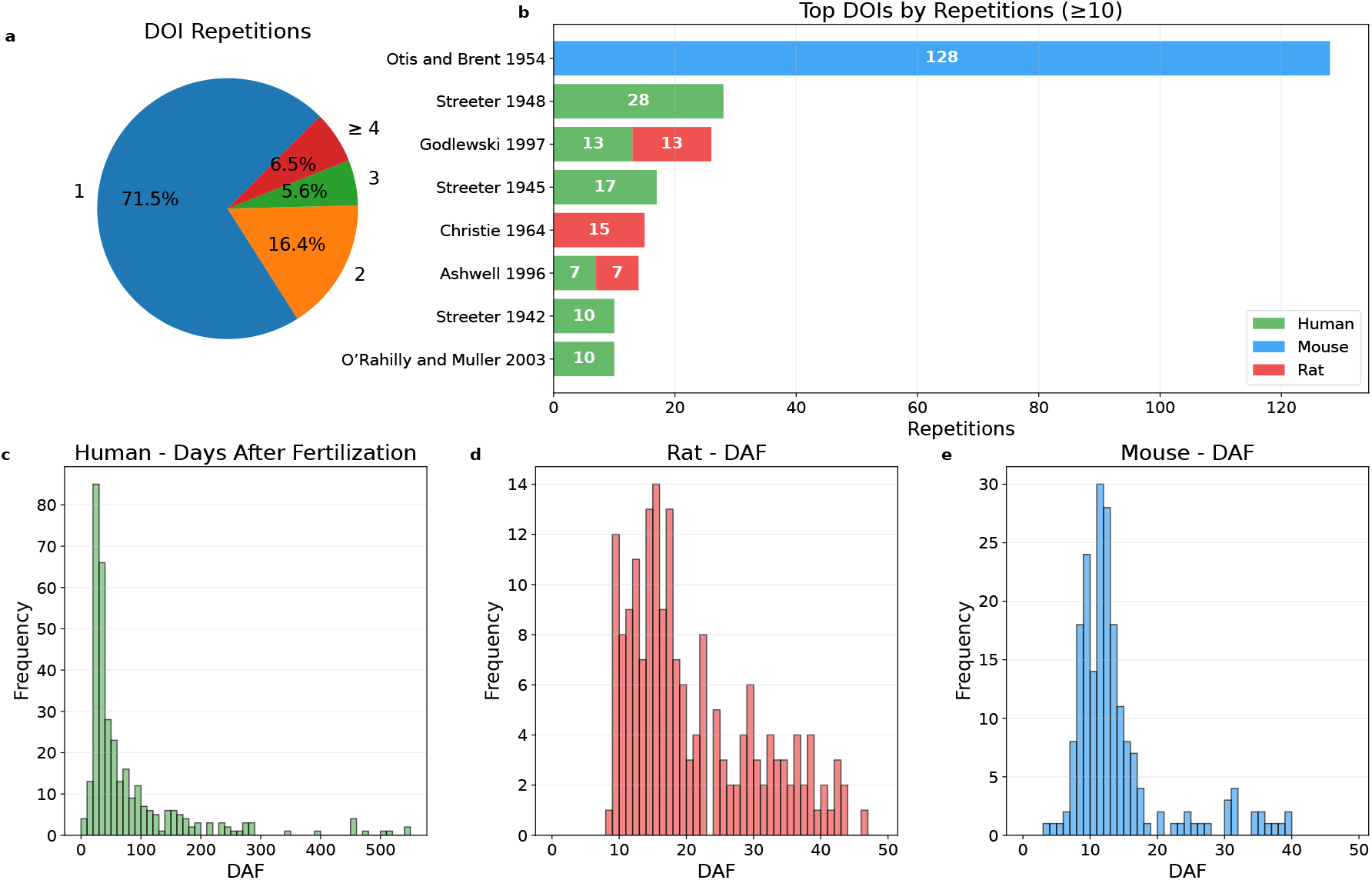
Analysis of source repetition and histograms of age density of analyzed milestones. (a) Distribution of DOI repetition frequency across the dataset, showing that most references occur only once, while a smaller proportion are repeated as sources of milestones. (b) Most frequently repeated DOIs (*≥* 10 repetitions), also showing the contribution of each species’ data for every reference. (c–e) Distribution of observations according to days after fertilization (DAF) for Human (c), Rat (d), and Mouse (e), illustrating the distinct developmental-age ranges and sampling densities across species.

However, examination of the most frequently repeated sources (*≥* 10 repetitions) (Figure 2b) reveals the presence of “anchor” or foundational studies contributing a significant number of milestones. The work of Otis and Brent (Otis and Brent 1954) stands out notably, contributing 128 mouse references to the dataset. Streeter (Streeter 1948) follows with 27 records and represents one of the main sources of human developmental observations incorporated into the work of Otis and Brent. Repeated contributions from other researchers are also represented in the dataset, although less frequently. This high repetition generates a notable concentration of mouse data primarily attributable to this classic work. To counteract this bias and enrich the variability of the dataset, data generated ad hoc for this study (Amici) account for approximately one-third of the total mouse data, thereby increasing the dataset’s variability and enhancing its generalizability.

Finally, the distribution of observations in DAF was analyzed for each species (Figure 2c-e). The histograms clearly illustrate the differences in temporal scales and variations in sampling density, reflecting the periods of organogenesis and prenatal development that have been historically more studied in each animal model.

### Human-rodent developmental relationships

Consistent with previous studies, the analysis of DAF patterns between humans and rodents revealed a nonlinear relationship. As shown in (Figure 3a and b), the overall dispersion of the data increases progressively with increasing DAF in both mice and rats, indicating that the variability associated with cross-species correspondence is not constant throughout development. To further illustrate the relationship during early development, a zoomed view of the data for DAF *≤* 300 is included (Figure 3c and d), highlighting the developmental period of particular relevance to the perinatal transition. This view shows that the same general non-linear pattern observed across the full range is maintained during early development. In other words, the relationships between human and rodent DAF are neither deterministic nor linear. These patterns were examined in greater depth in the companion work, including the role of an additive constant and the evaluation of alternative formulations to the logarithmic model widely used in the literature. The results indicated that developmental relationships were best described by a quarter-power model with an additive constant:

**Figure 3:**
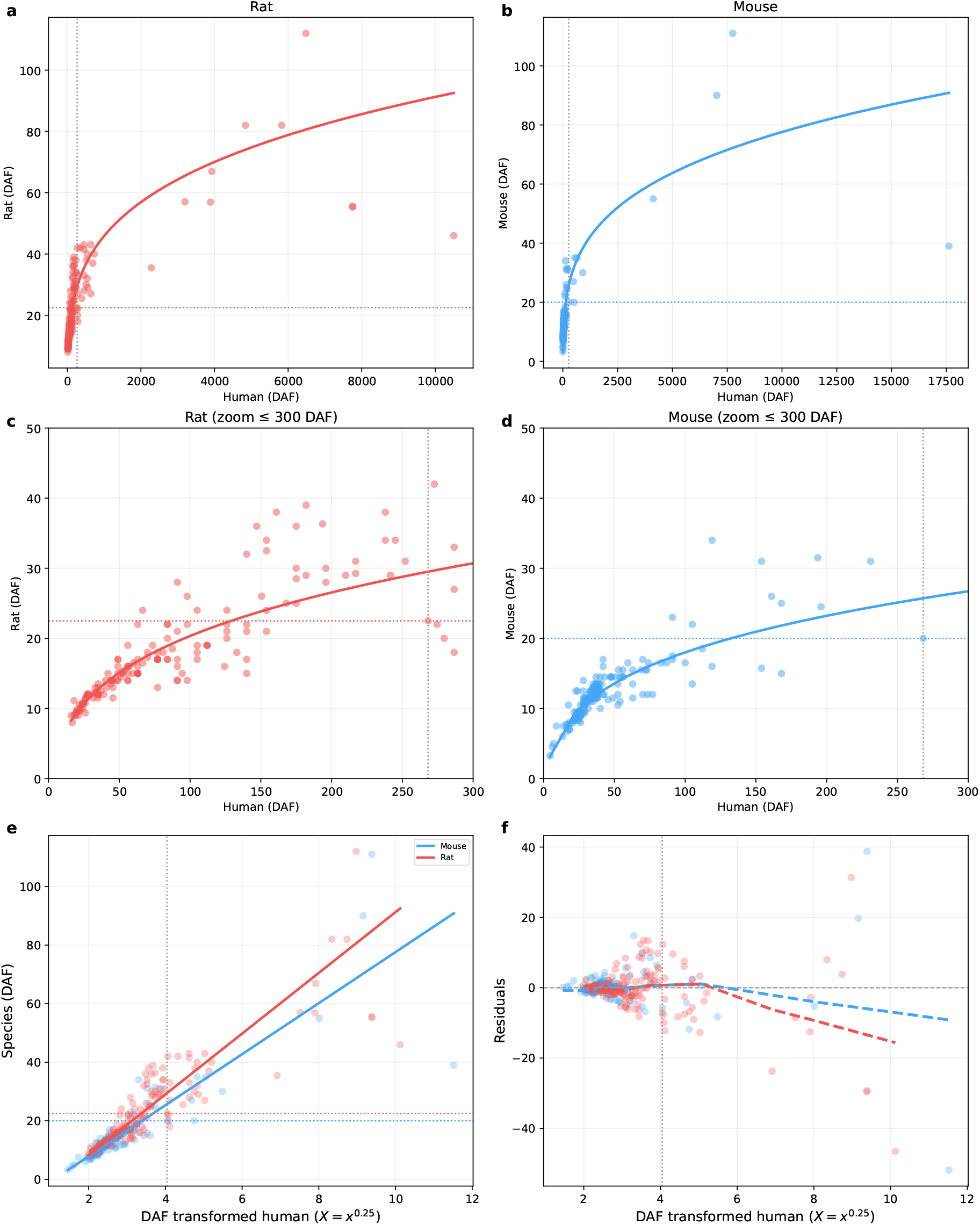
Distribution of rodent developmental milestones in DAF. **(a–b)** Scatter plots of Human DAF versus Rat and Mouse DAF, respectively, with WLS fits obtained in the companion study. **(c–d)** Same data restricted to Human DAF *≤* 300 to magnify the early-development region. **(e)** Linearity assessment after the *x*^0.25^ transformation (*X* = Human^0.25^), with weighted least-squares linear fits for Rat and Mouse. **(f)** Residuals from the linear WLS fits plotted against *X*, with LOESS-smoothed trend lines used to diagnose residual curvature. Dashed lines indicate the birth date for each species (red, rat; blue, mouse; black, human).

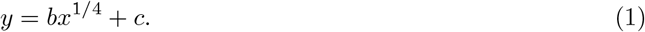

This model represents the nonlinear relationship between developmental timescales using a parsimonious function while incorporating a constant shift between the two scales. The corresponding transformation of the variables allows the relationship to be expressed in linear form, as shown in Figure 3e, where the transformed data exhibit a linear pattern. The parameter estimates from the companion work were *b* = 10.3716 and *c* = −12.4535 for rat, and *b* = 8.7156 and *c* = −9.5489 for mouse. These parameters were used to transform human DAF into the corresponding rodent developmental age, facilitating direct cross-species comparison.

Finally, model adequacy was examined through analysis of the residual distribution (Figure 3f). The residuals showed an approximately random dispersion across the range with the greatest density of observations, with no systematic curvature evident during the early developmental stages. This behavior indicates that the model does not exhibit substantial systematic deviations in the region with the greatest data support.

A deviation was nevertheless observed at the upper end of the DAF range. Although this deviation occurred in the same direction in both mice and rats, this region contains fewer observations and therefore entails greater uncertainty in the estimated trend. Consequently, the observed deviation may reflect a boundary effect associated with the limited availability of data during later developmental stages. However, its consistent occurrence in both species is noteworthy and could also indicate a systematic deviation from the model in this region, which should be evaluated using datasets with greater representation of advanced developmental stages.

### Rat-mouse developmental relationships

As mentioned above, the dataset contains a relatively small number of developmental milestones shared between mouse and rat. Despite the limitations imposed by this restricted overlap, a preliminary analysis was performed to assess the relationship between developmental ages in the two species. A simple linear regression model was fitted to the shared milestones, with the results shown in Figure 4. The assumptions of linear regression, including linearity, homoscedasticity, and normality of the residuals, were explored both analytically (Jarque-Bera *JB* = 2.677, *p* = 0.262; Breusch-Pagan *LM* = 2.889, *p* = 0.089)and visually. The analysis revealed a strong and statistically significant linear association between rat and mouse developmental ages (*R*^2^ = 0.909, *n* = 26). The estimated slope was 0.945 (95% CI: 0.819–1.070, *p <* 0.001), which was not significantly different from unity, while the intercept was not significantly different from zero (−0.829, 95% CI: −4.283–2.625, *p* = 0.625). Although the LOESS smoothing line shows slight curvature, the assumption of an overall linear relationship appears reasonable given the limited number of shared milestones. These results suggest that, within the range of available data, rat and mouse developmental ages are approximately linearly related, with a near one-to-one correspondence. This finding is consistent with the close evolutionary relationship and broadly similar developmental trajectories of the two species. However, the limited number of shared milestones warrants caution, as more extensive datasets may reveal more complex relationships. It is worth noting that the companion paper identified significant differences between rat and mouse developmental trajectories relative to humans.

**Figure 4:**
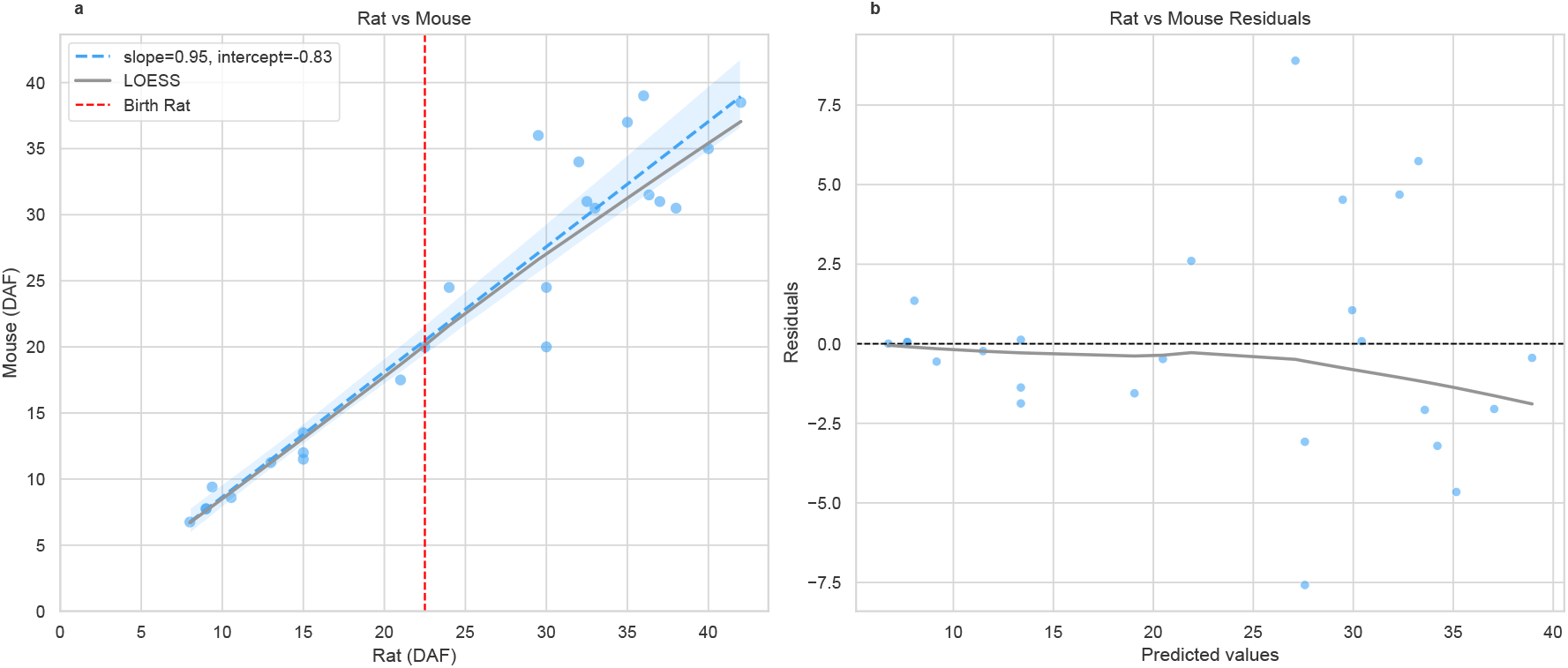
Rat–mouse developmental relationship. **(a)** Scatter plot of Mouse DAF as a function of Rat DAF, with the fitted linear regression (dashed line) and LOESS-smoothed trend (grey) shown for reference. The red dashed vertical line indicates rat birth. **(b)** Residuals from the linear fit plotted against fitted values, with a LOESS trend line (grey).

### Similar patterns of developmental scaling between brain and body milestones

To evaluate whether developmental scaling relative to humans differed between brain and body milestones, weighted linear fits relating linearized human age (*X* = Human^0.25^) to rodent DAF were compared between the Body and Brain tissue clusters, separately for each species (Figure 5). Within mice, the Body cluster (slope = 8.995) and Brain cluster (slope = 7.871) yielded broadly similar scaling relationships, and a joint F-test for simultaneous differences in slope and intercept did not reach significance (*F* = 1.95, *p* = 0.145). Neither the slope difference (∆_slope_ = −1.12, *p* = 0.133) nor the intercept difference (∆_intercept_ = 2.23, *p* = 0.197) was significant when considered individually. A parallel analysis in rats likewise showed no evidence of cluster-specific scaling (*F* = 0.48, *p* = 0.620; ∆_slope_ = −0.04, *p* = 0.959; ∆_intercept_ = −0.30, *p* = 0.886), with Body and Brain milestones following nearly identical trajectories (slopes of 10.55 and 10.51, respectively).

**Figure 5:**
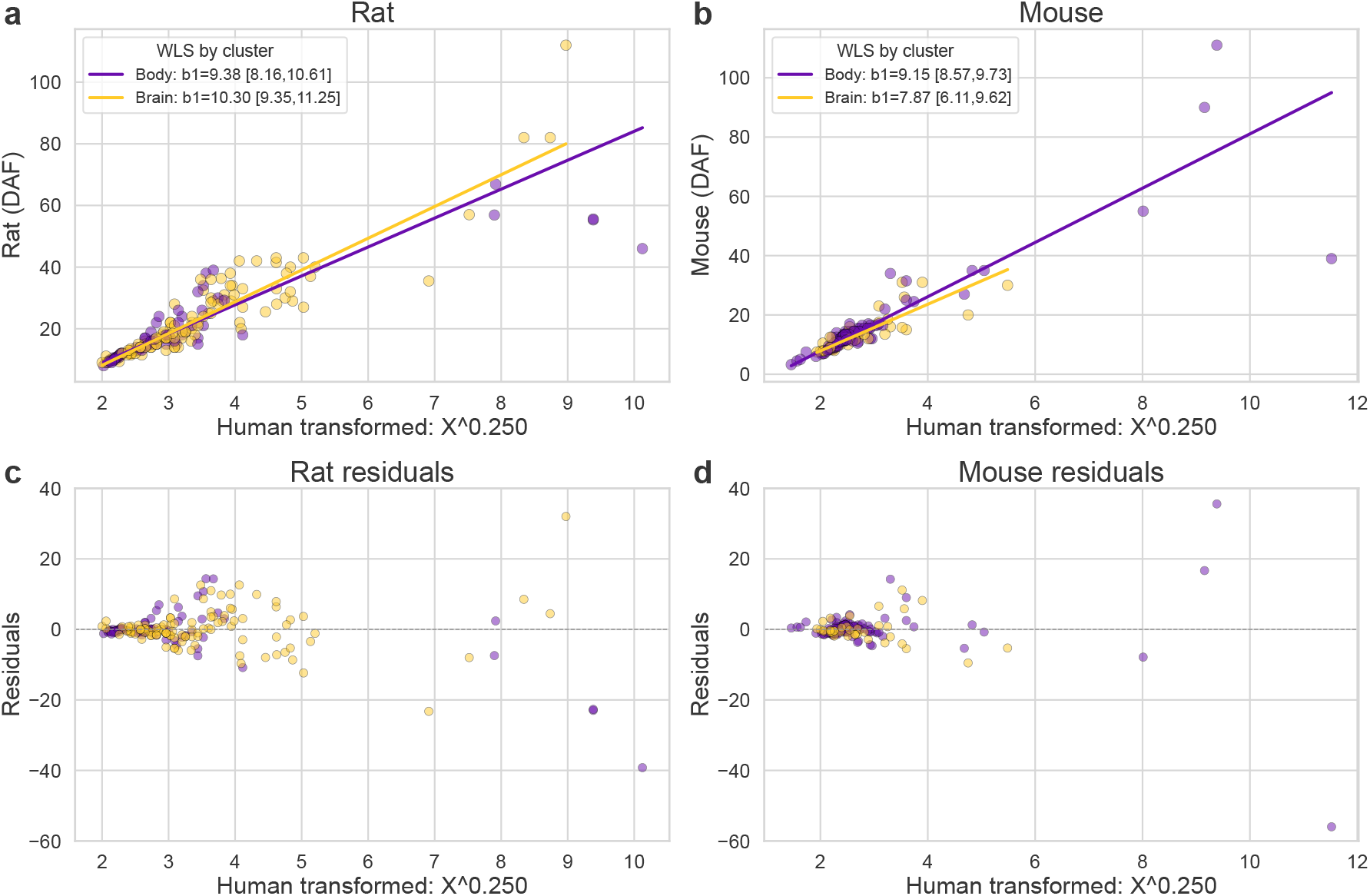
Allometric scaling of developmental age in humans, rats and mice, separated by tissue cluster. Human age was linearized as *X* = Human^0.25^. **(a**,**b)** Weighted least squares (WLS) fits of DAF as a function of linearized human age for Body (violet) and Brain (yellow) clusters in Rat (a) and Mouse (b). Points represent individual parameter values; lines represent WLS fits. Slope estimates and 95% confidence intervals are provided in the legend for each cluster. **(c**,**d)** Residuals from the corresponding WLS fits in (a) and (b), plotted against linearized human age, for Rat (c) and Mouse (d). Weights were derived from a variance model regressing log(residuals^2^) on log(*X*) to account for heteroscedasticity.

Finally, to assess whether the Body/Brain distinction contributed additional explanatory power beyond species identity, a cluster effect was tested within the joint rat + mouse model after controlling for species. This effect did not reach, conventional statistical significance (*F* = 2.50, *p* = 0.084), with neither the slope difference (∆_slope_ = −0.77, *p* = 0.138) nor the intercept difference (∆_intercept_ = 1.47, *p* = 0.242) attaining significance. Taken together, these results indicate that, once species identity is accounted for, the developmental scaling of Body and Brain milestones relative to humans is broadly comparable.

## Discussion

Here, we present a curated, open-access dataset of common neurodevelopmental and general developmental milestones for mice, rats, and humans. This resource expands upon our previous rat–human database (Campos Eusebi et al. 2024) by incorporating newly compiled mouse data. Consistent with earlier comparative work, the relationship between human and rodent developmental ages is distinctly non-linear. Furthermore, the present study shows that this relationship cannot be fully linearized using conventional logarithmic transformations. To identify a more robust framework, we tested allometric power-law models in a companion study. In that work, a quarter-power formulation with an additive constant emerged as the best-performing model. During the compilation of this dataset, we also generated a modest amount of overlapping developmental data for rats and mice. Interestingly, we observed a close correspondence between their developmental trajectories. Unlike the human–rodent relationship, when rat and mouse milestones were compared directly, they exhibited an approximately linear, near one-to-one relationship, with a slope not significantly different from unity and an intercept not significantly different from zero. This finding aligns with the close evolutionary relationship and broadly similar developmental programs of these two rodent species, and it contrasts sharply with the markedly non-linear pattern observed when either species is compared with humans. However, when rat and mouse trajectories were evaluated in terms of their scaling relative to human development, they differed significantly, in agreement with the species-specific parameters identified in the companion study. These observations raise a fundamental question: is quarter-power scaling applicable exclusively to human-rodent comparisons, or does it hold utility for analyzing developmental relationships across a broader range of species? This apparent discrepancy between the rodent–human relationship and the relationship between rodents themselves may stem from two underlying causes. First, mice and rats share a closer evolutionary history, having diverged only 12–24 million years ago, whereas the human and rodent lineages diverged approximately 75-90 million years ago (Rat Genome Sequencing Project Consortium et al. 2004). Furthermore, rats and mice have relatively similar body masses, particularly compared with humans, and share broadly comparable ecological and life-history strategies. Consequently, they may not have experienced the evolutionary constraints that typically give rise to pronounced power-law scaling. Second, if the non-linear effect is subtle, detecting it may require a substantially larger dataset, particularly given the overall biological similarity between these two species. Beyond the evidence presented in this work, it can be speculated that greater evolutionary distance may be associated with increased heterochrony, leading to greater divergence in developmental trajectories and, potentially, greater heteroscedasticity. Conversely, differences in developmental timescales may reflect variation in body mass, lifespan, and other life-history traits, which could be captured, at least in part, by the coefficients of a quarter-power scaling model (West et al. 2002).

### Modeling perinatal events

Another phenomenon evident in the dataset is the ontogenetic shift that occurs during the transition to extrauterine life, as rodents and humans experience this process at fundamentally different levels of developmental maturity. Mice and rats are born before attaining numerous milestones that humans have already achieved by birth, a pattern that is clearly reflected in our data. When the data are divided into four quadrants based on species-specific birth dates, a large cluster of milestones appears in the upper right quadrant; these represent events that occur postnatally in rodents but prenatally in humans.

In altricial species, the degree of somatic and neurophysiological maturity at birth is closely related to reproductive strategy and, in particular, to litter size. Polytocous reproduction involves a trade-off between the number of offspring and the maternal investment available to each individual, such that the production of large litters tends to be associated with smaller offspring and a lower degree of developmental maturity at birth (Charnov and Ernest 2006). This relationship is also integrated with interspecific differences in body size and production rates. Among mammals, larger-bodied species generally tend to produce smaller litters, whereas smaller species tend to produce larger litters and relatively smaller offspring (Tuomi 1980). This association is consistent with allometric constraints on production rate and resource allocation: as body size increases, metabolic and production rates per unit of body mass generally decrease, potentially limiting the amount of biomass that can be allocated to reproduction per unit of time. However, the relationships among body size, litter size, and neonatal maturity are unlikely to be driven exclusively by metabolic constraints. Rather, they reflect the evolutionary integration of multiple life-history traits, including gestation length, offspring size, mortality, and ecological conditions (Sibly and Brown 2009). Within this framework, the relatively advanced developmental state of human neonates compared with those of highly polytocous rodents, such as mice and rats, may reflect broader interspecific differences in reproductive strategies, prenatal investment, and the physical constraints associated with carrying a larger offspring. The longer human gestation relative to that of mice and rats may be related not only to greater prenatal investment per offspring, but also to biomechanical constraints on maternal reproduction that limit the size and developmental state of the offspring that can be carried to term. This phenomenon has fueled considerable debate in the literature regarding the applicability of neonatal animal models to human perinatal pathologies. Our stance aligns with previous work (Ortiz et al. 2022): developmental relationships should be understood as ranges rather than one-to-one correspondences. Consequently, when modeling developmental processes, researchers must evaluate the physiological processes affecting their specific model and consider the potential impact of developmental offsets in other processes. This dataset aims to serve as a tool to facilitate such evaluations. In this context, relying on postnatal development in rodents to reach a developmental stage comparable to that of humans may not always be an appropriate approach.

Another source of debate in the literature has been tissue-specific considerations in comparative development, such as the brain growth spurt hypothesis, among others (Dobbing and Sands 1979). Here, comparisons of body and brain milestone clusters revealed no significant differences. Furthermore, the observed differences were not consistent across the two rodent species. These findings suggest that developmental scaling relative to humans does not differ substantially between neural and non-neural milestones. Thus, brain development appears to follow a developmental trajectory that is broadly coordinated with overall somatic development. This partially contrasts with the popular anthropocentric view that humans possess a superior, or at least exceptional, brain. At least in terms of the overall pattern of brain development, however, our results do not indicate a clear departure from the developmental patterns observed in other mammals. To explore this less anthropocentric perspective of evolution further, see (Striedter et al. 2012). Furthermore, an important question in biology is whether development and aging can be considered two sides of the same biological process (Silva-García 2023). Substantial evidence indicates that development, growth, body size, reproduction, aging, and longevity are closely interrelated, a concept sometimes referred to in the literature as the “pace of life” (Yuan et al. 2023). Within this context, species exhibiting faster development and maturation tend to display higher reproductive investment (i.e., more offspring) and accelerated aging, whereas those with a slower pace of life show the opposite trajectory. This dataset supports this perspective, as it provides a unified model for early, mid, and late development. Given that the dataset is open-access, it could be integrated in the future with comparative aging data to further investigate this topic.

### Limitations and future directions

Several limitations should be noted. First, a small number of highly cited bibliographic sources contribute disproportionately to specific portions of the mouse dataset. This reduces variability, potentially affecting the generalizability of the models (Vázquez-Borsetti 2024). For example, Otis and Brent (1954) (Otis and Brent 1954) is an excellent source of data, with almost no exclusions based on our inclusion criteria. However, heavy reliance on it poses a generalization problem because the authors conducted all their studies using a single mouse strain. This also introduces uniform breeding conditions, homogeneous handling, and consistent researcher criteria, thereby artificially reducing variability. While Otis is the most prominent example, other sources with high repetition exist, which may limit the independence of some estimates. Efforts were made to include additional data to diversify sources and improve generalizability; thus, the dataset can be used as is. However, it is advisable to verify whether the analyses hold when data from these dominant authors are excluded. Naturally, the future inclusion of more milestones will help mitigate this issue. The second limitation arises in the comparative analysis of rats versus mice. The number of milestones shared between the two species remains relatively small, warranting caution when generalizing the observed linear correspondence. Larger, more balanced datasets could reveal more complex or non-linear relationships. Finally, the retrospective and literature-based nature of the dataset means that reported ages inherit the methodological heterogeneity of the original studies. While this heterogeneity can paradoxically improve generalizability by capturing diverse laboratory conditions, it also introduces noise; therefore, standardization, strict inclusion criteria, and curation procedures were applied to mitigate this issue.

Overall, this open dataset and the associated analyses provide a transparent and extensible resource for translating developmental timing across mouse, rat, and human, complementing the modeling framework developed in the companion paper. Continued community contributions and the inclusion of additional milestones, particularly those shared across all three species, should further improve the precision and generalizability of cross-species developmental translation.

## AI use statement

The dataset was compiled and curated entirely manually, without the use of artificial intelligence tools for data extraction or curation. AI-assisted tools were used to support code development and to improve the clarity and language of the manuscript. All AI-assisted outputs were reviewed and validated by the authors, who take full responsibility for the content of the dataset, the analyses, and the manuscript.

## Data Availability

The dataset generated and analyzed in this study, together with the code used for data processing, statistical analyses, and figure generation, is openly available in the GitHub repository https://github.com/Vazquez-Borsetti/mouse-rat-human-development. The dataset is additionally permanently archived with a citable DOI in Zenodo: https://doi.org/10.5281/zenodo.23244347, and the code is permanently archived with its own citable DOI in Zenodo: https://doi.org/10.5281/zenodo.23246067. The repositories contain the data used in the analyses and the code required to reproduce the reported results.

## References

Bergsjø, Per et al. (Jan. 1990). “Duration Of Human Singleton Pregnancy: A Population-based Study”. In: Acta Obstetricia et Gynecologica Scandinavica 69.3, pp. 197–207. issn: 0001-6349, 1600-0412. doi: 10.3109/00016349009028681. (Visited on 09/01/2026).

Bryda, Elizabeth C. (2013). “The Mighty Mouse: The Impact of Rodents on Advances in Biomedical Research”. In: Missouri Medicine 110.3, pp. 207–211. issn: 0026-6620.

Campos Eusebi, Wanda et al. (Feb. 2024). “Divergent Pattern of Development in Rats and Humans”. In: Neurotoxicity Research 42.1, p. 7. issn: 1029-8428. doi: 10.1007/s12640-023-00683-y.

Charnov, Eric L. and S. K. Morgan Ernest (Apr. 2006). “The Offspring-Size/Clutch-Size Trade-Off in Mammals”. In: The American Naturalist 167.4, pp. 578–582. issn: 0003-0147, 1537-5323. doi: 10.1086/501141. (Visited on 09/24/2026).

Clancy, B., R. B. Darlington, and B. L. Finlay (2001). “Translating Developmental Time across Mammalian Species”. In: Neuroscience 105.1, pp. 7–17. issn: 03064522. doi: 10.1016/S0306-4522(01)00171-3.

Dewar, A. D. (Apr. 1968). “LITTER SIZE AND THE DURATION OF PREGNANCY IN MICE”. In: Quarterly Journal of Experimental Physiology and Cognate Medical Sciences 53.2, pp. 155–161. issn: 0033-5541. doi: 10.1113/expphysiol.1968.sp001954. (Visited on 09/04/2026).

Dobbing, John and Jean Sands (Mar. 1979). “Comparative Aspects of the Brain Growth Spurt”. In: Early Human Development 3.1, pp. 79–83. issn: 03783782. doi: 10.1016/0378-3782(79)90022-7. PMID: 118862.

Donaldson, H. H. (Sept. 1918). “A Comparison of Growth Changes in the Nervous System of the Rat with Corresponding Changes in the Nervous System of Man”. In: Proceedings of the National Academy of Sciences 4.9, pp. 280–283. issn: 0027-8424. doi: 10.1073/pnas.4.9.280.

Gjerdevik, M. et al. (May 2026). “Gestational Age Determination in Pregnancies Conceived via Assisted Reproductive Technology”. In: Ultrasound in Obstetrics & Gynecology 67.5, pp. 604–612. issn: 0960-7692, 1469-0705. doi: 10.1002/uog.70219. (Visited on 09/04/2026).

Hain, A. M. (Apr. 1936). “THE PHYSIOLOGY OF PREGNANCY IN THE RAT: FURTHER DATA ON THE PASSAGE OF HORMONES VIA THE PLACENTA AND THE MOTHER’S MILK”. In: Quarterly Journal of Experimental Physiology 26.1, pp. 29–43. issn: 0370-2901. doi: 10.1113/expphysiol.1936.sp000699. (Visited on 09/04/2026).

Ortiz, Mauro, Fabián Loidl, and Pablo Vázquez-Borsetti (Sept. 2022). “Transition to Extrauterine Life and the Modeling of Perinatal Asphyxia in Rats”. In: WIREs Mechanisms of Disease 14.5. issn: 2692-9368. doi: 10.1002/wsbm.1568.

Otis, Eileen M. and Robert Brent (1954). “Equivalent Ages in Mouse and Human Embryos”. In: The Anatomical Record 120.1, pp. 33–63. issn: 10970185. doi: 10.1002/ar.1091200104.

Rat Genome Sequencing Project Consortium et al. (Apr. 2004). “Genome Sequence of the Brown Norway Rat Yields Insights into Mammalian Evolution”. In: Nature 428.6982, pp. 493–521. issn: 0028-0836, 1476-4687. doi: 10.1038/nature02426. (Visited on 09/09/2026).

Romijn, Herms J., Michel A. Hofman, and Albert Gramsbergen (1991). “At What Age Is the Developing Cerebral Cortex of the Rat Comparable to That of the Full-Term Newborn Human Baby?” In: Early Human Development 26.1, pp. 61–67. issn: 03783782. doi: 10.1016/0378-3782(91)90044-4.

Sibly, Richard M. and James H. Brown (June 2009). “Mammal Reproductive Strategies Driven by Offspring Mortality-Size Relationships”. In: The American Naturalist 173.6, E185–E199. issn: 0003-0147, 1537-5323. doi: 10.1086/598680. (Visited on 09/24/2026).

Silva-García Carlos Giovanni (May 2023). “Devo-Aging: Intersections Between Development and Aging”. In: GeroScience 45.4, pp. 2145–2159. issn: 2509-2723. doi: 10.1007/s11357-023-00809-2. (Visited on 09/10/2026).

Streeter, George L. (1948). “Contributions to Embryology.” In: (visited on 10/07/2026).

Striedter, Georg F., John C. Avise, and Francisco J. Ayala (June 2012). “In the Light of Evolution VI: Brain and Behavior”. In: Proceedings of the National Academy of Sciences 109.Supplement_1, pp. 10607–10611. issn: 0027-8424, 1091-6490. doi: 10.1073/pnas.1206166109. (Visited on 09/10/2026).

Tuomi, Juha (Apr. 1980). “Mammalian Reproductive Strategies: A Generalized Relation of Litter Size to Body Size”. In: Oecologia 45.1, pp. 39–44. issn: 0029-8549, 1432-1939. doi: 10.1007/BF00346705. (Visited on 09/24/2026).

Vázquez-Borsetti, Pablo (Dec. 2024). “Variability in Rat Weight Gain during Development”. In: Laboratory Animals 58.6, pp. 579–590. issn: 0023-6772, 1758-1117. doi: 10.1177/00236772241246370. (Visited on 09/10/2026).

West, Geoffrey B., William H. Woodruff, and James H. Brown (Feb. 2002). “Allometric Scaling of Metabolic Rate from Molecules and Mitochondria to Cells and Mammals”. In: Proceedings of the National Academy of Sciences 99.Suppl_1, pp. 2473–2478. issn: 0027-8424, 1091-6490. doi: 10.1073/pnas.012579799. (Visited on 09/10/2026).

Yuan, Rong et al. (Nov. 2023). “Relationships among Development, Growth, Body Size, Reproduction, Aging, and Longevity – Trade-Offs and Pace-Of-Life”. In: Biochemistry (Moscow) 88.11, pp. 1692–1703. issn: 0006-2979, 1608-3040. doi: 10.1134/S0006297923110020. (Visited on 08/18/2026).

